# Mechanism-based classification of SARS-CoV-2 Variants by Molecular Dynamics Resembles Phylogenetic Tree

**DOI:** 10.1101/2023.11.28.568639

**Authors:** Thais Arns, Aymeric Fouquier d’Hérouël, Patrick May, Alexandre Tkatchenko, Alexander Skupin

## Abstract

The COVID-19 pandemics has demonstrated the vulnerability of our societies to viral infectious disease. The mitigation of COVID-19 was complicated by the emergence of Variants of Concern (VOCs) with varying properties including increased transmissibility and immune evasion. Traditional population sequencing proved to be slow and not conducive for timely action. To tackle this challenge, we introduce the Persistence Score (PS) that assesses the pandemic potential of VOCs based on molecular dynamics of the interactions between the SARS-CoV-2 Receptor Binding Domain (RBD) and the ACE2 residues. Our mechanism-based classification approach successfully grouped VOCs into clinically relevant subgroups with higher sensitivity than classical affinity estimations and allows for risk assessment of hypothetical new VOCs. The PS-based interaction analysis across VOCs resembled the phylogenetic tree of SARS-Cov-2 demonstrating its predictive relevance for pandemic preparedness. Thus, PS allows for early detection of a variant’s pandemic potential, and an early risk evaluation for data-driven policymaking.

## Introduction

Since the emergence of COVID-19 in Wuhan, China in late 2019, the disease has significantly impacted global health [Wang, 2022] with over 767 million confirmed cases and approximately 6.9 million deaths as of November 2023 [WHO, 2023]. COVID-19, caused by the SARS-CoV-2 virus, lead to atypical viral pneumonia [Wu & McGoogan, 2020] with an immune response similar to SARS and MERS. The virus spread quickly worldwide [Deng, 2020], and despite the development of vaccines and treatments, it continued to challenge public health systems and demonstrates the vulnerability of our modern societies to viral infectious diseases. Non-pharmaceutical interventions have been required to prevent healthcare systems from being overwhelmed. Variants of the virus, such as Alpha, Delta, and Omicron, have contributed to surges in cases due to increased transmissibility [Bushman, 2021; Liu, 2021; Planas, 2021] and reduced vaccine effectiveness [Grabowski, 2021]. The Omicron variant, reported in November 2021, was particularly concerning due to its 51 mutations in the spike protein and its ability to partially evade immunity. However, its milder symptoms and lower hospitalization rates, especially among vaccinated individuals [Callaway, 2021], have led to a relaxation of the pandemic severeness and represent a step towards endemics, however the effect of future VOCs can be only barely estimated. While population sequencing allows to identify VOCs by their increasing prevalence only with a significant delay, mitigation strategies would benefit from an early assessment of potential risks from new virus variants.

Transmissibility of the SARS-CoV-2 virus is strongly linked to the densely glycosylated transmembrane Spike (S) proteins protruding from the viral surface to enter human cells [Barros, 2021]. The S protein is a trimeric fusion protein that consists of subunits, S1 and S2. S exists in a meta-stable pre-fusion conformation, which undergoes a substantial structural rearrangement when binding the host cell membrane receptor [Li, 2016]. Structurally it presents flexibility that translates into an ensemble of angiotensin-converting enzyme 2 (ACE2) homodimer conformations that could sterically accommodate binding of the S protein trimer to more than one ACE2 homodimer and suggests a mechanical contribution of the host receptor toward the large S protein conformational changes required for cell fusion [Barros, 2021]. This process is triggered when the S1 subunit binds to a host cell’s ACE2 type I membrane protein. The receptor binding proceeds through docking of the receptor-binding domain (RBD) of the viral S protein to the peptidase domain (PD) of ACE2 and destabilizes the pre-fusion trimer resulting in shedding of the S1 subunit and transition of the S2 subunit to a stable post-fusion conformation [Walls, 2017]. The RBD is a 211 amino acid region (residues 319–529) at the C-terminus of S1, which is essential for virus entry and the presumed target of neutralizing antibodies [Shang, 2020]. Hence, it plays a central role in increased transmissibility and reduced vaccine efficacy [Burioni, 2021, Piccoli, 2020].

Since late 2020, various VOCs of the SARS-CoV-2 virus have emerged with convergent amino acid substitutions (**Table 1**). The N501Y substitution is present in the Alpha, Beta, Gamma, and Omicron variants, and increases the virus’s binding affinity to ACE2 receptors [Starr, 2020]. The E484K substitution is found in Alpha2, Beta, and Gamma variants and has been associated with the virus’s ability to evade the immune response from monoclonal antibodies and antibodies in convalescent plasma [Weisblum, 2020; Greaney, 2021]. The Beta, Delta2, Gamma, and Omicron variants have additional substitutions K417N and K417T [Wise, 2021]. Mutations L452R and T478K are associated with the Delta variant, with K417N observed in a sub-lineage called Delta2 [Tao, 2021]. The K417 substitutions have lesser impact on polyclonal antibody responses compared to substitutions like E484K [Greaney, 2021; Barnes, 2020]. These substitutions are also expected to slightly reduce the virus’s binding affinity to ACE2 receptor [Starr, 2020]. The Gamma variant, characterized by K417T, E484K, and N501Y substitutions, is estimated to have 1.7 to 2.4 times higher transmissibility, and prior infections provide 54% to 79% protection against this variant [Faria, 2021].

**Table 1.** Considered SARS-CoV-2 VOCs (including the official Pango lineage [Rambaut, 2020] and respective mutated residues.

| Variant | Mutated Residues |
| --- | --- |
| **Alpha** (B.1.1.7) | N501Y |
| **Alpha2** (B.1.1.7+E484K) | E484K, N501Y |
| **Beta** (B.1.351) | N501Y, K417N, E484K |
| **Delta** (B.1.617.2) | L452R, T478K |
| **Delta2** (B.1.617.2.2) | L452R, T478K, K417N |
| **Gamma** (B.1.1.28.1) | N501Y, E484K, K417T |
| **Omicron** (B.1.1.529.1) | G339D, S371L, S373P, S375F, K417N, N440K, G446S, S477N, T478K, E484A, Q493R, G496S, Q498R, N501Y, Y505H |
| **Omicron BA.2** (B.1.1.529.2) | G339D, S371F, S373P, S375F, T376A, D405N, R408S, K417N, N440K, S477N, T478K, E484A, Q493R, Q498R, N501Y, Y505H |
| **Omicron BA.3** (B.1.1.529.3) | G339D, S477N, T478K, E484A, Q493R, Q498R, N501Y, Y505H |
| **Omicron BA.4** (B.1.1.529.4) | G339D, S371F, S373P, S375F, T376A, D405N, R408S, K417N, N440K, L452R, S477N, T478K, E484A, F486V, Q498R, N501Y, Y505H |
| **Omicron BA.5** (B.1.1.529.5) | G339D, S371F, S373P, S375F, T376A, D405N, R408S, K417N, N440K, L452R, S477N, T478K, E484A, F486V, Q498R, N501Y, Y505H |
| **Omicron BA.2.12.1** (B.1.1.529.2.12.1) | G339D, S371F, S373P, S375F, T376A, D405N, R408S, K417N, N440K, L452Q, S477N, T478K, E484A, Q493R, Q498R, N501Y, Y505H |
| **Deltacron** (AY.4 + BA.1) | G339D, S371L, S373P, S375F, K417N, N440K, G446S, S477N, T478K, E484A, Q493R, G496S, Q498R, N501Y, Y505H, |
| **Omni** | G339D, S371L, S373P, S375F, T376A, D405N, R408S, K417N, N440K, G446S, L452R, S477N, T478K, E484K, F486V, Q493R, G496S, Q498R, N501Y, Y505H |
| **P2** (B.1.1.28.2) | E484K |

The Omicron BA.1 variant of SARS-CoV-2 has 51 missense amino acid mutations, with 32 located in the S protein, including 15 substitutions in the receptor binding domain (RBD) that interacts with host ACE2 receptors and is a major target of neutralizing antibodies. This shows significantly more mutations in the RBD compared to the Alpha, Beta, Gamma, and Delta variants, which have 1, 3, 3, and 2 mutations in the RBD, respectively [EU/EEA, 2021]. The numerous mutations in the RBD of the Omicron variant could affect its infectivity, transmissibility, and the efficacy of vaccines and therapeutic antibodies [Liu, 2021; Cao, 2021; Callaway, 2021]. Studies showed that the Omicron variant had an increased risk of reinfection compared to primary infection [Pulliam, 2021]. The variant also spread rapidly, with a doubling time of 3.18–3.61 days, outcompeting the Delta variant and becoming the dominant strain globally [Grabowski, 2021]. Neutralizing antibody responses to Omicron were reduced compared to the original virus and Delta variant in vaccinated individuals, but booster doses enhanced antibody levels [Cele, 2021; Wilhelm, 2021]. The Omicron variant showed lower severity, with 65% lower risk of hospitalization or death and 83% lower risk of ICU admission or death compared to Delta, though the high transmissibility of Omicron could still strain healthcare systems [Ulloa, 2022]. Protection from previous infection or vaccination and intrinsically reduced virulence of the Omicron variant contributed to the lower severity, with an estimated 25% reduced risk of severe hospitalization or death compared to Delta [Davies, 2022]. Omicron variant has derivative lineages, including BA.2 to BA.5. The WHO reported that BA.5 represented over half of the current global cases, while BA.4 accounts for just over 10% [WHO Weekly epidemiological update on COVID-19, 2022]. The spread of BA.5 highlights the unpredictable nature of the pandemic and the potential for new Variants of Concern (VOCs) to cause significant epidemic rebounds.

While these insights emphasize the central role of the S protein for the pandemic dynamics and highlight the importance of specific mutations, as well as the interactions between proteins as the main drivers for biological processes, a more systematic understanding allowing for a more reliable variant classification is still elusive. Here, we describe the Persistence Score (PS), a new method to evaluate the risk potential in terms of increased transmissibility of virus variants that can be assessed by molecular dynamics investigations of the viral S protein considering the contact and/or loss of contact between SARS-CoV-2 RBD and ACE2 residues, outperforming classical energy-based (ι1G) approach and inferred couplings between putatively interacting residues, revealing that the PS-based interaction analysis across VOCs resembled the phylogenetic tree. The highly detailed molecular data is subsequently used as a measure of molecular interaction at the mutation site providing a risk assessment also for potential future recombinant variants like Deltacron allowing for early adaptation of mitigation strategies of political decision makers.

## Material and Methods

### Molecular modeling

To classify SARS-CoV-2 variants based on the interaction interfaces of the ACE2 and RBD proteins, we applied a full-atom molecular dynamics (MD) simulations approach. The sequence similarity of ACE2 and RBD between SARS-CoV-2 and SARS-CoV is only 73% [Andersen, 2020], precluding existing models for studying SARS-CoV-2 VOCs. Several 3D structures are available in the Protein Data Bank (PDB) [Berman, 2000] for the detailed study of the S protein of SARS-CoV-2, such as 6VXX (closed-state conformation) and 6VYB (open-state conformation), but these massive structures contain more than 1280 amino acid residues with low experimental resolution, several gaps in the structure and missing residues. Since most mutations of concern are concentrated in the interface between ACE2 and RBD, we focused on a high-resolution crystallography model as reference template for the VOC modeling, the WT SARS-CoV-2 structure (PDB ID 6LZG [Wang, 2020]), which provided the most accurate molecular interaction data. All structures are available in the **Supplementary Material** and GitLab (https://git-r3lab.uni.lu/ICS-lcsb/ercsacov/).

### Molecular dynamics (MD) simulations

MD simulations were performed in triplicates, for a total of 600 ns for each of the SARS-CoV-2 variants using GROMACS v2020 [Lindahl, 2020] and CHARMM36 force field [Huang, 2017]. A cubic box was defined with at least 9 Å of liquid layer around the protein, using single-point charge water model and periodic boundary conditions. An appropriate number of sodium (Na^+^) and chloride (Cl^−^) counter-ions were added to neutralize the system at the final concentration of 0.15 mol/L. The algorithms V-rescale (*τ_t_* = 0.1 ps) and Parrinello-Rahman (*τ_p_* = 2 ps) were used for temperature and pressure coupling, respectively. Cut-off values of 1.2 nm were used for both van der Waals and Coulomb interactions, with Fast Particle-Mesh Ewald (PME) electrostatics. For all MD simulations, the production stage was preceded by three steps of Energy Minimization (alternating steepest-descent and conjugate gradient algorithms), and eight steps of equilibration as previously described [Devaurs, 2017, Arns, 2020]. Briefly, the Equilibration stage started with position restraints for all heavy atoms (5,000 kJ^−1^mol^−1^nm^−1^) and a temperature of 310 K, for a period of 300 ps, to allow for the formation of solvation layers. The temperature was then reduced to 280 K and the position restraints were gradually reduced. This process was followed by a gradual increase in temperature (up to 300 K). Together, these equilibration steps represented the first 500 ps of each simulation. During the production stage, the system was held at constant temperature (310 K) without restraints. The Cα Root Mean Square Deviation (RMSD) and Root Mean Square Fluctuations (RMSF) values were calculated using the initial structures as reference.

### Historical sequences (Mock controls)

To demonstrate that the PS filtering and inferred couplings clustering are not biased or related to the chosen methodology, we created mock controls from historical sequences (randomly generated mocks and early SARS-CoV-2 mutations) as reported on GISAID [Khare, 2021]. The considered mock mutations were Mock_Free_01 K386E, D398S, R457A; Mock_Free_02: K356N, E465Q, C480F; Mock_Free_03: D405I, V511D, H519T; Mock_Weighted_01: F338L, G476S, S438F; Mock_Weighted_02: A522S, Q414E, V367F; Mock_Weighted_03: A520S, S494P, N439K. The nomenclature for the amino acid residue changes for the Historical sequences and Omni variant is as follows: K356N = original amino acid residue (K), mutation position (356) and mutated amino acid residue (N).

### Omni Variant (synthetic variant)

To assess the impact of all Omicron-related mutations, a synthetic variant named *Omni* was modeled, which included all Omicron mutations considered in this study (Omicron, Omicron BA.2, Omicron BA.2.12.1, Omicron BA.3, Omicron BA.4, Omicron BA.5): G339D, S371L, S373P, S375F, T376A, D405N, R408S, K417N, N440K, G446S, L452R, S477N, T478K, E484K, F486V, Q493R, G496S, Q498R, N501Y, Y505H.

### Persistence Score

Based on the raw MD data, we established a Persistence Score (PS) that considers the contact and/or loss of contact between SARS-CoV-2 RBD and ACE2 residues, which is subsequently used as a measure of molecular interaction and to assess levels of three-dimensional (3D) structural deformation at the mutation site or in the vicinity. The PS is calculated based on the molecular interactions observed during the MD simulations by PS = (interaction time x 100) / (simulation duration) and provides an estimate of spatially resolved binding and may therefore be indicative of transmissibility. Interaction time was calculated using PyMol 2.4.2 [Schrödinger, 2015] with a default distance threshold of 1 Å between interacting residues.

### Free Energy Calculations

Free energy calculations were performed using the *gmx_MMPBSA* [Valdés-Tresanco, 2021] package and respective GROMACS v2020 [Lindahl, 2020] trajectory files. The binding free energy *(11G_bind_)* of the RBD-ACE2 complex system were obtained by the following equation:

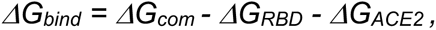

where *11G_com_*, *11G_RBD_*, and *11G_ACE2_* were the free energies of the complex, RBD and ACE2, respectively. For each system, 20 frames were extracted from the 200 ns trajectory for *ΔG* calculation. Total binding free energies using the ACE2 and RBD proteins were calculated for all variants and replicates, as well as the per residue decomposition schemes.

### Inferred couplings between putatively interacting residues

Atomic coordinates from MD data were extracted using *gmx dump -f* (GROMACS v2020 [Lindahl, 2020]) and reported positions were averaged by residue, yielding coordinate matrices *X_n,t_*, *Y_n,t_*, and *X_n,t_* for residue *n* at time-point *t*. From these coordinate matrices, a residue root-mean-square (RMS) matrix was computed as

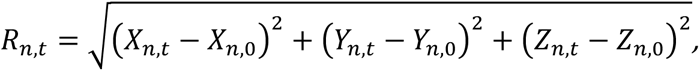

which allowed for the fast evaluation of RMSD and RMSF by summing over residues and time-points, respectively. Couplings between residues were inferred using the Thouless-Anderson-Palmer (TAP) approximation of the solution to the inverse Ising problem [Nguyen, 2017]. The inferred coupling between residues *i* and *j* are thus given as

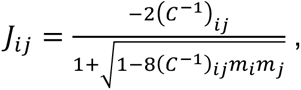

where *C* is the covariance matrix between residue positions *m*_&_ and *m*_’_as the average positions of the residues across the simulation’s timeframe. Since the covariance matrix is generally not uniquely invertible, its inverse is computed as the Moore-Penrose generalized inverse (*ginv* function of package MASS version 7.3-54 in GNU R version 4.04), which can lead to numerical instabilities. Clustering of simulated variants was performed by averaging inferred couplings across replica and considering residue ranges, which participate in the direct interaction between ACE2 and RBD, specifically between ACE2(19,49), ACE2(61,87), ACE2(322,330), ACE2(351,357), ACE2(383,393) and RBD(403,408), RBD(417,421), RBD(437,458), RBD(473,506), and RBD(610,620), respectively.

### Principal Component Analysis

PCAs were performed using *PCAtools* R package [Blighe and Lun; 2019] with R version 4.2.2 (2022-10-31).

### VOC distances from MD simulations

For Euclidean distances, the corresponding rotated PCA values for each variant were subsequently used to calculate the Euclidean distance between variants. The relationship between VOCs was further characterized by the dendrograms obtained from the clustering of the three interaction analysis considering PS, affinity and coupling estimations. For this purpose, the resulting dendrograms were characterized by the distance measures of the *tree* package in R in analogy to the phylogenetic distance.

### Phylogenetic distances

The phylogenetic data was sourced from the NextStrain [Hadfield, 2018] platform by downloading the Nexus tree file containing the SARS-CoV-2 relevant data (based on nucleotide sequences), which was parsed and read into R version 4.2.2 (2022-10-31) using the *read.nexus* function from the *ape* package. The Euclidean distance between the branches of the phylogenetic tree was calculated using the *cophenetic.phylo* function from the *ape* package, resulting in a matrix of distances. The distance matrix obtained was converted to a dendrogram object, providing a visual representation of the phylogenetic relationships among the SARS-CoV-2 variants, with branch lengths representing the Euclidean distances.

## Results

### Phylogeny and structural flexibility of the ACE2-RBD interactions based on variant specific mutations

The dynamics of the COVID-19 pandemic was driven by the appearance of VOCs as shown by the timeline of the NextStrain phylogenetic data of variants starting with the Alpha variant by the end of 2020, followed by the Beta, Gamma, Delta and the Omicron subfamily (**Fig. 1A**). Each variant came with its specific set of mutations in the RBD affecting the 3D structure of the ACE2 and RBD interaction regions (**Fig. 1B**). Molecular dynamics simulations of the considered VOCs **(Table 1)** indicated common flexible regions throughout the entire ACE2 protein structure by the normalized Root Mean Square Fluctuation (RMSF) (**Fig. 1C**). Interestingly, around residues 300 – 320, low to slightly negative values were found for some of the variants, such as for Delta2 and Omicron, while P2 and Alpha had values larger than 0.1 Å. The normalized RMSF of the RBD (**Fig. 1D**) showed an unstable area for all Omicron variants around residues 370 – 380, whereas the instability around residues 380 – 400 was specific for the Omicron BA.3, Gamma and Deltacron variants. The region around residues 440 – 460 showed clear RMSF peaks for the Beta and Alpha2 variants, while the region of residues 475 – 490 showed a unique 0.2 Å normalized RMSF peak for Alpha2 and Omicron BA.5.

**Figure 1.**
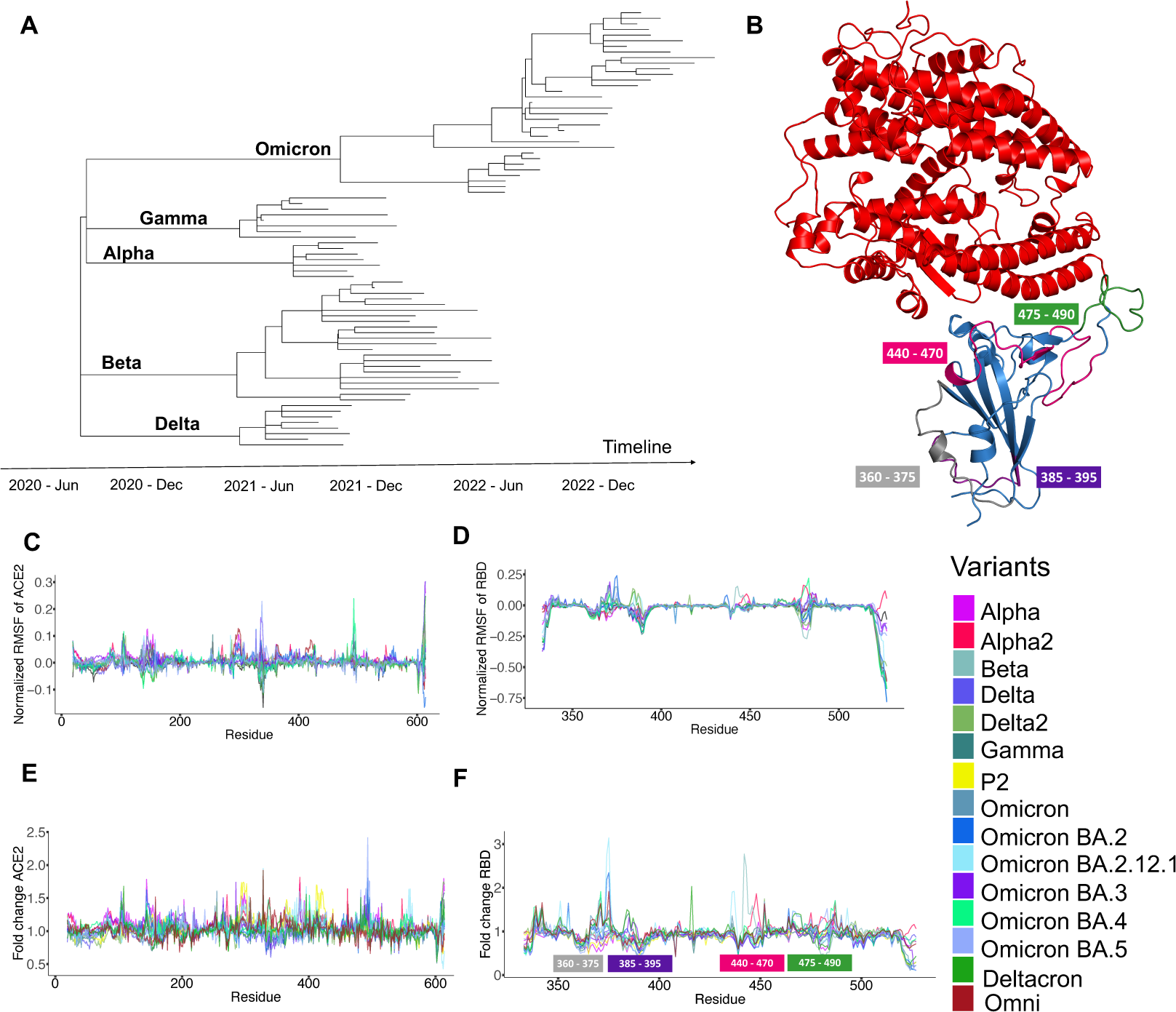
SARS-CoV-2 evolution and structural analysis of the RBD highlights flexible regions. **(A)** Timeline of SARS-CoV-2 variants. **(B)** SARS-CoV-2 WT model (**ACE2**: red; **RBD**: blue) depicting specific areas of interest (also shown in **(F)**). **(C)** and **(D)** depict the normalized RMSF for ACE2 and RBD, respectively. **(E)** and **(F)** depict the fold change for ACE2 and RBD, respectively.

To further classify the observed structural flexibility, we analyzed the variant specific fold change of the RMSF normalized to the WT strain for the ACE2 (**Fig. 1E**) and RBD (**Fig. 1F**) interfaces. For ACE2, the structural changes spread over the entire structure, while the instabilities within the RBD were localized in specific protein segments as shown by the structural location in the color-coded structure (**Fig. 1B**). All analyzed variants displayed a very low Root Mean Square Deviation (RMSD) (2 to 5 Å), indicating that all variants retain their 3D structure flexible, but without major secondary structure changes when considering the ACE2-RBD structure **(Supplementary Fig. 1A)**. Further analysis showed that ACE2 exhibited an almost perfect superimposition in RMSF for all SARS-CoV-2 variants (**Supplementary Fig. 1B**) whereas the RBD exhibited variant-specific levels of structural flexibility for several amino acid regions (**Supplementary Fig. 1C**) with the most notably regions for residues 360 to 375, 385 to 395, 440 to 470 (specifically for variants Alpha2 and Beta) and 475 to 490.

### Persistence Score classifies mutation-induced changes in ACE2-RBD binding in a structure-dependent manner

To investigate whether the changes in flexibility has an impact on the interaction between the RBD and the ACE2 in VOC specific manner and can be used for classification, we developed and applied the Persistence Score (PS) as a sensitive measure of binding activity (**Methods**) of the virus variants (**Fig. 2**). The analysis of the 3 independent simulations for each considered VOC identified 41 interacting residues for ACE2 (**Fig. 2A**). The nomenclature for the amino acid residue changes considering the protein chain, amino acid and residue position is as follows: BTYR501 = chain B, residue TYR and position 501. The resulting PS signature sorted by decreasing values in the WT strain exhibits strain specific differences on top of a trend for consistently high PS values (>90) and thus persistent interactions for all variants in positions 30 (AASP30), 24 (AGLN24), 34 (AHSD34), 31 (ALYS31), 353 (ALYS353), 28 (APHE28), 27 (ATHR27), 41 (ATYR41), 83 (ATYR83), and 355 (AASP355). Compared to the WT strain, residue 82 (AMET82) exhibited lower PS values for Omicron BA.2, Omicron BA.4, Omicron BA.5 and the synthetic variant Omni. Residue 19 (ASER19) presented a high PS in the WT (>90), which dropped for the Alpha, Alpha2, Beta, P2, Delta, Delta2 and Gamma variants (12, lowest PS). As for the Omicron subvariants, Omicron BA.3, Omicron BA.4 and Omicron BA.5 showed the lowest PS (∼70) for residue 19. Residue 393 (AARG393) showed PS (∼50) for WT, while Alpha, Alpha2, P2 and Delta had higher PS values (>90), and considerably lower PS for Omicron (∼20), Omicron BA.2.12.1 (15), and similar values for Omicron BA.2, Omicron BA.3, Omicron BA.4, Omicron BA.5 and Deltacron (∼30). In addition to these main differences, VOC specific interactions were also associated with residues 20 (ATHR20) (high in variants Delta, Delta2 and Gamma), 75 (AGLU75) (higher PS for Beta and Gamma variants) and 356 (APHE356) (highest PS for Omicron BA.2.12.1). For the RBD (**Fig. 2B**), 50 interacting residues were identified with a trend for consistently high (>90) PS for all variants in positions 475 (BTYR495), 493 (BARG493/BGLN493), 498 (BARG498/BGLN498), 505 (BHSD505/BTRY505), 455 (BLEU455), 456 (BPHE456), 486 (BPHE486/BVAL486), 500 (BTHR500), 453 (BTRY453), 473 (BTYR473), 501 (BASN501/BTYR501) and 502 (BGLY502). The PS obtained for position 417 (BASN417/BLYS417/BTHR417) was the highest (100) for Alpha, Alpha2, P2, Delta and Omicron BA.3 and reduced for Beta (90), Delta2 (90), Gamma/Omicron BA.2 (50), Omicron (75), Omicron BA.2.12.1 (80), Omicron BA.4, Omicron BA.5 and Deltacron (70). Residue 484 (BALA484/BGLU484/BLYS484) exhibited an increased PS for the Beta variant (80) compared to the WT strain (60) and variants Alpha, Alpha2, P2 and Delta, while all other variants had values lower than 50. Additional discriminating residues were 495 (BTYR495) with high PS (>80) for the Gamma and most Omicron variants compared to Deltacron (16), and residue 483 (BARG493/BGL493) with highest PS for the Alpha variant (50), followed by Omicron BA.4 and Omicron BA.5 (40).

**Figure 2.**
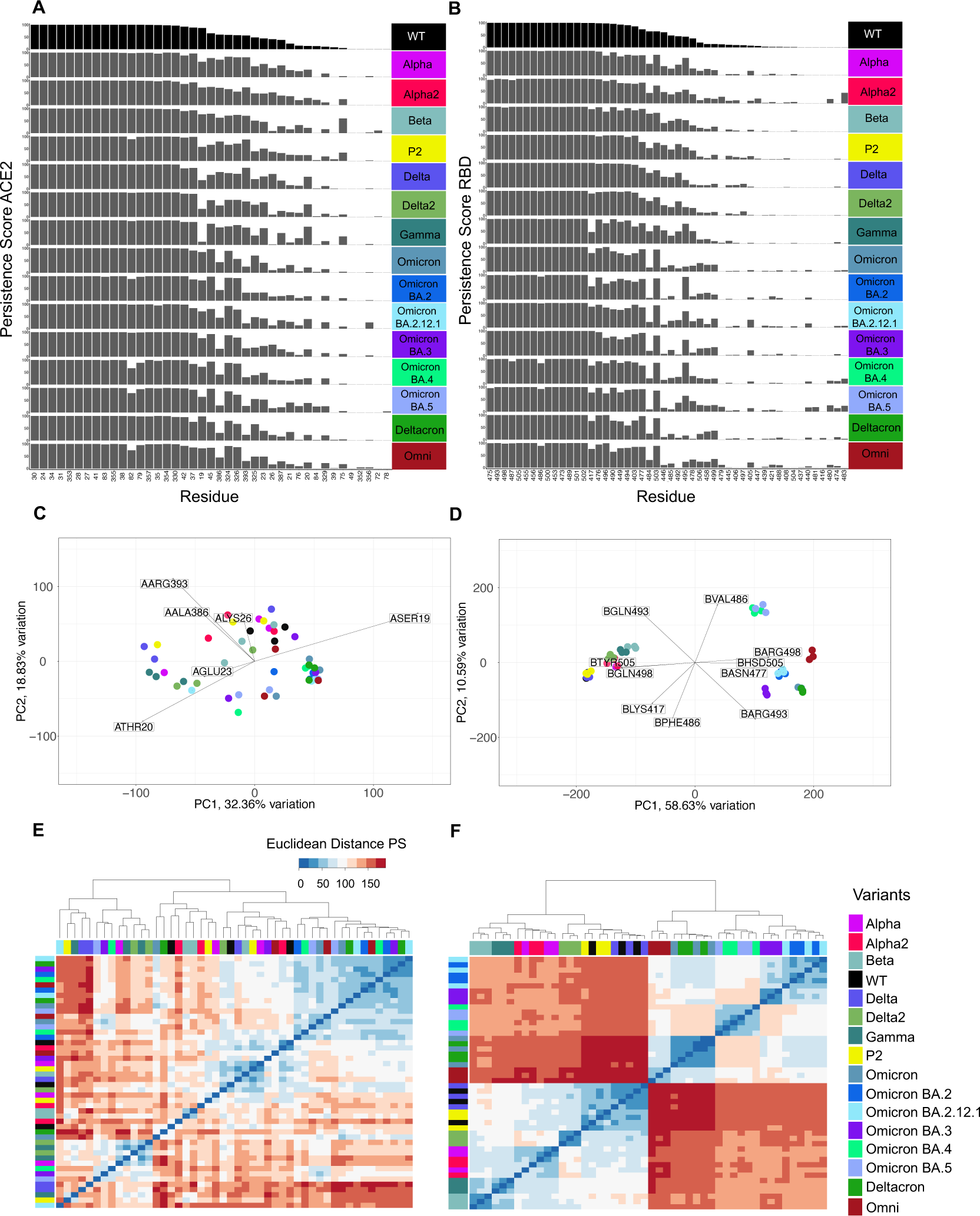
Persistence score allows for VOC classification. **(A)** Persistence score for ACE2 residues. **(B)** Persistence score for RBD residues. **(C)** PCA of ACE2 for all considered SARS-CoV-2 variants. **(D)** PCA of RBD for all considered SARS-CoV-2 variants. (**E)** Euclidean distance clustering for ACE2. **(F)** Euclidean distance clustering for RBD.

To investigate if the higher sensitivity of the PS allows for more robust grouping of VOCs, we performed Principal Component Analysis (PCA) to the residue-resolved binding signatures. For ACE2 the resulting PCA biplot (**Fig. 2C**) indicates a similar amount of explained variability (50.6%) and a slightly more structured pattern based on the main discriminating residues ASER19, ATHR20, AGLU23, ALYS26, AALA386, AARG393 compared to the affinity analysis (**Fig. 3C**). However, individual strains form again mixed groups with no clear pattern. By contrast, the PCA of the RBD (**Fig. 2D**) exhibits an increased amount of explained variability of 69% for the first 2 PC and clear grouping of VOC-specific realizations. The first PC separates the VOCs in two main groups based on the discriminating factors BGLN49, BTYR505, BARG498, BHSD505 and BASN477 into a group containing the Alpha, Alpha2, Beta, Gamma, Delta, Delta2, WT and P2 variants on the left, and the Omicron subvariants, the Deltacron and the synthetic Omni variant group on the right. Interestingly, the Omicron variants form 5 clearly separated subgroups *i)* Omicron and Deltacron; *ii)* Omicron BA.2 and Omicron BA.2.12.1; *iii)* Omicron BA.3; *iv)* Omicron BA.4 and Omicron BA.5; *v)* Omni along the 2^nd^ PC determined by the changed residues BPHE486/BVAL486 and BGLN493/BARG493. The clear separation into subgroups indicates the potential for VOCs classification by residue-resolved interactions analysis by PS. Interestingly, the Deltacron variant clusters together with the Omicron variant what may indicate that the recombinant would be more determined by the Omicron than the Delta variant properties. Furthermore, the synthetic Omni variant carrying all VOCs mutations does not exhibit a distinct behavior, but rather similar differences like between the individual Omicron subvariants.

**Figure 3.**
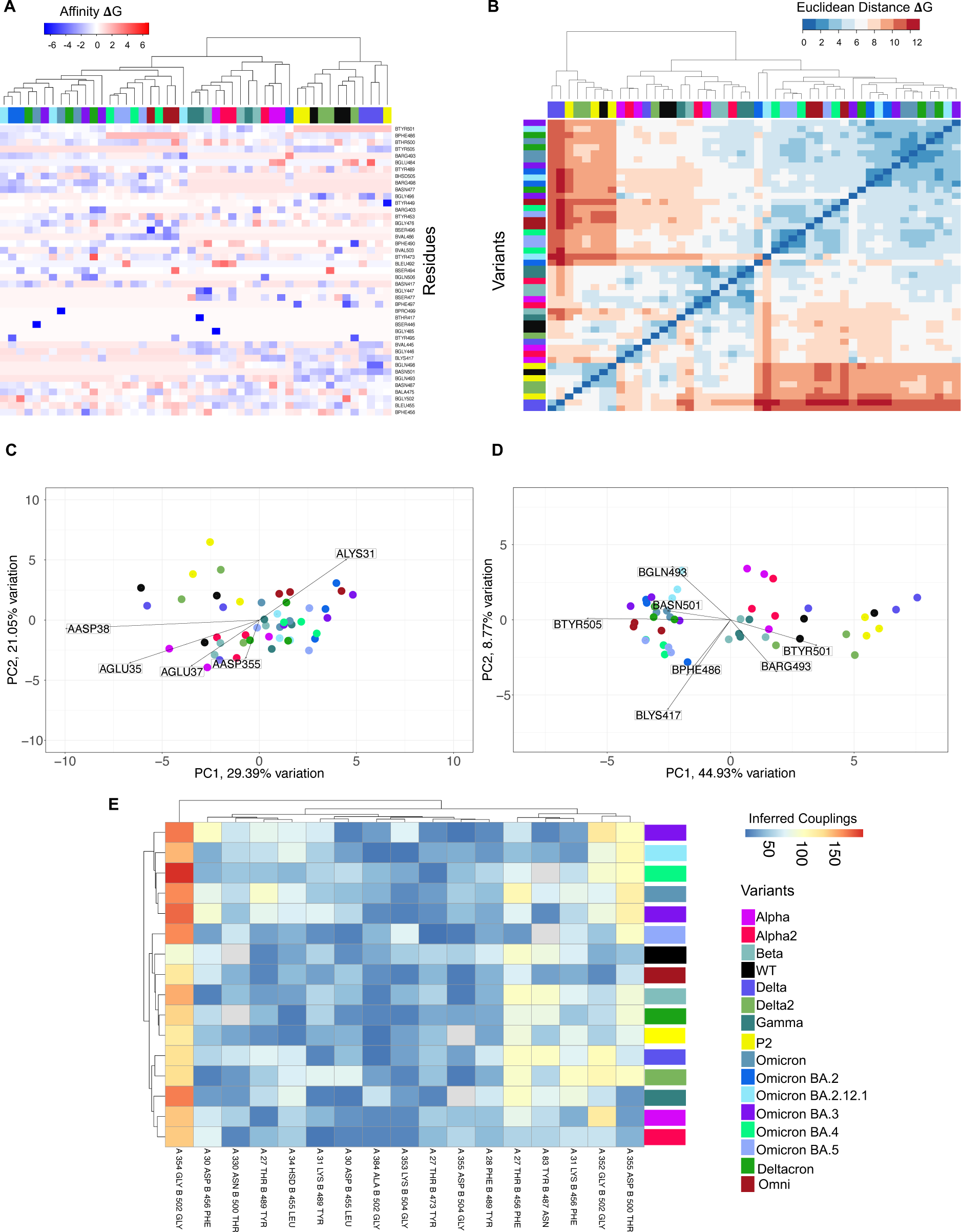
Interaction analysis of ACE2 and RBD for VOC strains indicates RBD significance. **(A)** Affinity ΔG heatmap, considering the RBD chain. **(B)** Euclidean distance of the residue ΔG values of the RBD. **(C)** PCA based on ΔG analysis of the ACE2 for all SARS-CoV-2 variants. **(D)** PCA based on ΔG analysis of the RBD for SARS-CoV-2 variants. **(E)** Strongest inferred coupling between ACE2 and RBD residues.

Based on the clearer clustering, we next calculated the Euclidean distance of the variants in the PS space and performed clustering for the ACE2 and the RBD interactions, respectively. The ACE2 interaction analysis did not cluster replicates and related variants into related subgroups (**Fig. 2E**). However, clustering of strains based on the RBD PS analysis, led to 2 big clusters where one contained the Omicron subvariants (Omicron BA.1, Omicron BA.2, Omicron BA.2.12.1, Omicron BA.3, Omicron BA.4, Omicron BA.5), as well as the Deltacron and Omni variants and the other group gathered the Alpha, Alpha2, Beta, Gamma, Delta, Delta2, P2 and WT variants (**Fig. 2F**). More detailed analysis revealed that replicates of individual strains closely related and related VOCs are typically grouped together like the Alpha and Alpha2 or Beta and Gamma variants.

### Variant classification using ΔG free energy binding

Given the VOC-specific interaction patterns identified by PS, we next tested how the binding free energies ΔG are affected by the VOC-specific mutations. The calculated affinities of the SARS-CoV-2 RBD-ACE2 complex **(Table 2)** exhibits the strongest binding energy of -66.20 kcal/mol for the P2 variant, while the WT complex showed the weakest binding energy of -40.83 kcal/mol. Interestingly, the estimated affinities did not revealed a clear pattern of strain relation where e.g. the Beta (-63.49 kcal/mol) and Omicron BA.3 (-63.31 kcal/mol) variants exhibited very similar values but are associated with rather different transmissibilities. Also, the affinities of the different Omicron variants exhibited rather different values which were not distinguishable from other VOCs. Similarly, the 2 Delta variants have rather different values (Delta: -60.17 kcal/mol vs Delta2: -47.31 kcal/mol) indicating that the overall affinity is not able to discriminate between variants.

**Table 2.** Considered SARS-CoV-2 variants and respective free energy binding values (ΔG_bind_).

| Variant | $\Delta G_{\text{bind}}$ |
| --- | --- |
| **Alpha** | -49.25 |
| **Alpha2** | -58.16 |
| **Beta** | -63.49 |
| **Control** | -40.83 |
| **Delta** | -60.17 |
| **Delta2** | -47.31 |
| **Gamma** | -43.43 |
| **P2** | -66.20 |
| **Omicron** | -61.88 |
| **Omicron_BA2** | -44.64 |
| **Omicron_BA3** | -63.31 |
| **Omicron_BA4** | -52.18 |
| **Omicron_BA5** | -46.56 |
| **Omicron_BA2121** | -57.32 |
| **Deltacron** | -57.41 |
| **Omni** | -54.19 |

To investigate whether the structural instabilities induce a residue dependent affinity pattern in a variant specific manner, we calculated the ΔG free energy per residue by energy decomposition. The resulting ΔG affinity heatmap for the RBD chain exhibits energies between -6 to +6 kcal/mol and hierarchical clustering grouped the variants in 3 main clusters (**Fig. 3A**). Most discriminating residues were BTYR501 which exhibited weaker binding energies for the P2, Delta, Delta2 and the WT variants compared to the other variants and BTYR505, which exhibits positive ΔG energies for all Omicron variants, the synthetic Deltacron and Omni strains, while the other variants have all negative ΔG energies. For residue BPHE486, the variants Omicron BA.4, Omicron BA.5 and Omni displayed weaker binding energies compared to all other variants whereas for residue BASN501 the variants P2, Delta, Delta2 and WT represented negative ΔG energies.

We next calculated the Euclidean distance within the ΔG space of the RBD chain to assess the potential to group variants into meaningful subgroups (**Fig. 3B**). The analysis shows that some variants cluster together within the same group with the largest distances for variants P2, Delta and Delta2 to the other variants, however the overall cluster composition exhibits a rather heterogenous picture with mixed variants (**Fig. 3B**) compared to the PS analysis (**Fig.2F**).

To further investigate the potential of residue resolved ΔG free energy for strain classification, we performed again PCA for the ACE2 (**Fig. 3C**) and for the RBD (**Fig. 3D**) profiles. The ACE2 analysis indicates the residues ALYS31, AGLU35, AGLU37, AASP38 and AASP355 as discriminating factors with around 50% of explained variability for the first 2 PC but individual realizations of the different variants do not show a clear pattern (**Fig. 3C**). The PCA of the RBD (**Fig. 3D**) exhibits a similar amount of explained variability and a separation between the Omicron subvariants, Deltacron and Omni, and the other variants (Alpha, Alpha2, Beta, Control, Delta, Delta2, Gamma, P2) based on residues BTYR505 and BTYR501. Despite this global separation, the different separations of the individual strains do not form strong individual clusters in the RBD PCA space compared to the PS analysis (**Fig.2D**) and has thus a more limited classification potential. Taken together, these results demonstrate that the PS of the RBD is more sensitive and superior tool to reveal residue interactions and allows for VOC classification, contrary to ΔG free energy binding approach.

### Inferred couplings between ACE2 and RBD allow for SARS-CoV-2 variant grouping

Given the limited classification power of the affinity analysis, we next investigated whether the interactions between specific residues of the RBD and ACE2 receptor can improve the grouping of VOCs. For this purpose, we inferred the couplings between the ACE2 and RBD residues by Thouless-Anderson-Palmer approximation (**Methods**). In the context of inferring couplings between putatively interacting residues, the Thouless-Anderson-Palmer (TAP) approximation of the solution to the inverse Ising problem [Thouless, 1977; Nguyen, 2017] is a sophisticated computational method to infer the strength and nature of interactions that we applied here between amino acid residues in proteins based on their correlated movements as observed in molecular dynamics simulations. This approach provided here a detailed understanding of protein interactions at a molecular level. The most significant couplings were subsequently used for clustering of variants and interactions (**Fig. 3E**). The clustering revealed interesting variant subgroups, such as the Omicron subvariants (Omicron BA.1, Omicron BA.2, Omicron BA.2.12.1, Omicron BA.3, Omicron BA.4), followed by a group containing the Omicron BA.5, WT, Omni, Beta, Deltacron, and P2 variants, and a group with the Delta, Delta2, Gamma, Alpha, and Alpha2 VOCs. The analysis also indicated the most significant couplings between the interactive residues, such as AGLY354/BGLY502 which had the highest scores for the Omicron subgroup (Omicron BA.4, Omicron BA.2, Omicron BA.3, Omicron BA.5, Omicron, Omicron BA.2.12.1) and Gamma variants, followed by the Beta lineage. Compared to the affinity analysis (**Fig. 3B**), the inferred couplings seem to reflect the epidemic relations between the VOCs better, but the separation of the original Omicron strain form the other Omicron variants as well as the grouping of the Beta and Gamma variants challenges a robust classification.

### PS groups VOCs in a pandemic relevant manner

To compare the 3 different classification approaches, the Euclidean distance between the variants was calculated for each classification space (PS, 11G free energy binding, inferred couplings). Subsequently, the distance was used for clustering and resulting dendrograms were analyzed (**Figs. 4A, B, C**). For the PS we observed a clear separation of the variants in the following subgroups (Delta2, P2, WT, Delta); (Alpha, Alpha2); (Beta, Gamma); (Omicron BA.4, Omicron BA.5; Omicron BA.3); (Omicron BA.2, Omicron BA.2.1.2.1); (Omni; Omicron, Deltacron). The clustering of the residue-resolved 11G free energy binding led to the groups (Omicron BA.2, Omicron BA.2.1.2.1); (Omicron BA.4, Omicron BA.5; Omicron BA.3; Omicron; Deltacron, Omni; Delta2, Delta, P2); (Alpha, Gamma); (WT, Alpha2, Beta). From the inferred couplings, we obtained the groups (Omicron BA.3, Omicron BA.2.1.2.1, Omicron BA.4, Omicron, Omicron BA.2); (Omicron BA.5, WT, Omni, Beta, Deltacron, P2); (Delta, Delta 2, Gamma, Alpha, Alpha 2). While all 3 approaches were able to separate most Omicron subvariants from the other VOCs, the subgrouping exhibited some differences between the approaches. Thus, PS grouped the 2 Alpha variants as well as the Beta and Gamma variants together, whereas the affinity-based clustering put the Alpha and Gamma variant together and grouped Alpha2 with the Beta and WT variants. In the inferred coupling analysis, the Omicron BA.5 variant is grouped together with the WT, Beta and P2 variants, in contrast to the 2 other approaches which group all Omicron related variants together in one major cluster. Thus, the PS approach seems to reflect the relations between the VOCs in a more pandemic relevant manner than the affinity and coupling based approaches. Taken together, these comparisons demonstrate that the PS of the RBD is more sensitive than the ΔG free energy binding and Inferred couplings to reveal residue interactions and allows for VOC classification.

**Figure 4.**
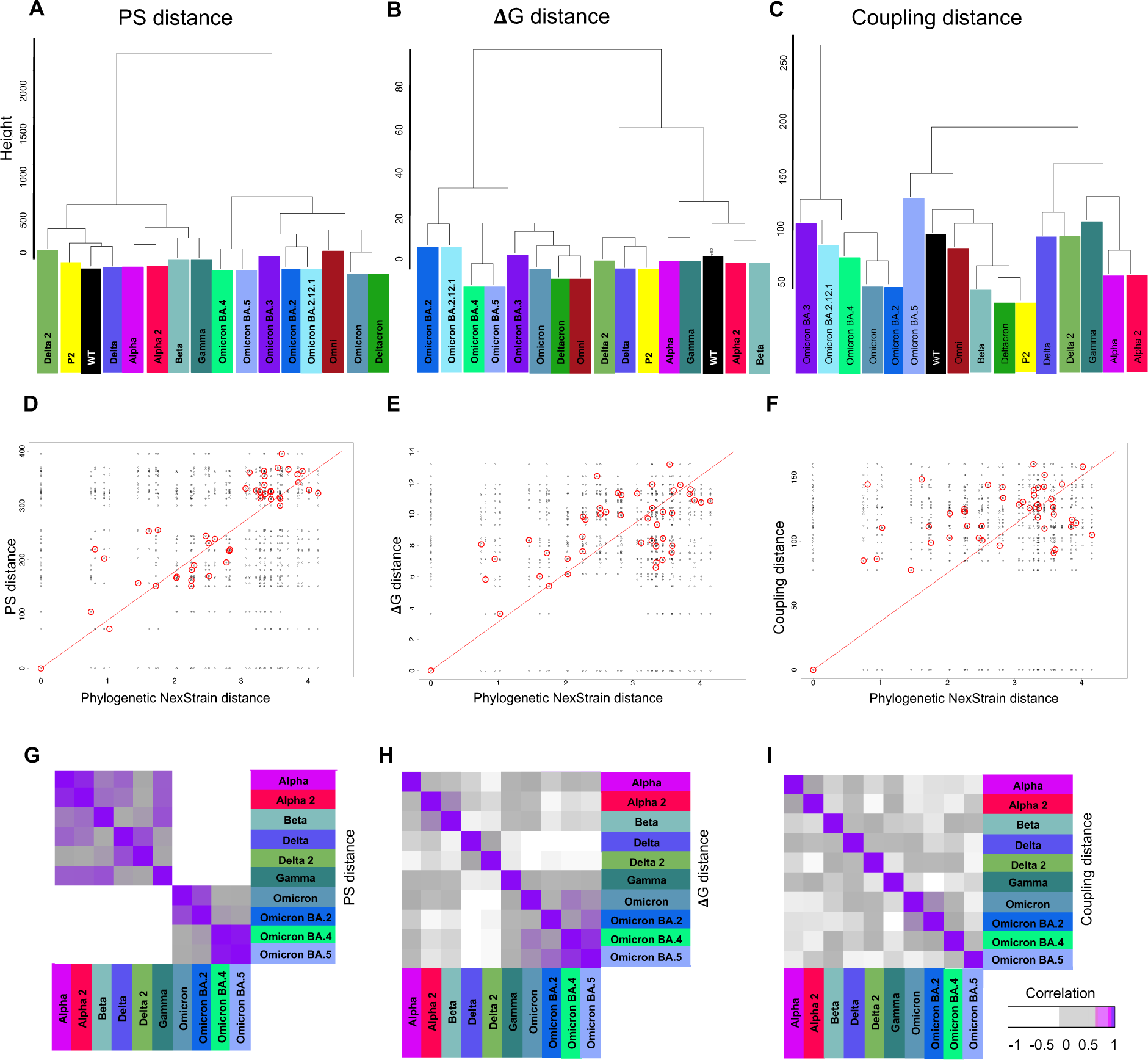
Persistence score based characterization resembles phylogenetic distance better than ΔG and inferred couplings. **(A-C)** Euclidean distance between the SARS-CoV-2 variants according to the applied methodology (PS, ΔG and Inferred coupling). **(D-F)** Distance analysis between phylogenetic Nexstrain distance and PS, ΔG and Coupling based distances, respectively. **(G-F)** Corresponding strain specificity of distance analyses.

### PS clustering resembles NextStrain-based phylogenetic tree

For a quantitative assessment of the obtained VOC grouping, we finally compared the interaction-based clustering with the phylogenetic information of NextStrain. For SARS-CoV-2, Nextstrain’s phylogenetic analyses and distance measurements are based on nucleotide sequences, which are aligned to a reference sequence. Nextstrain uses this aligned sequence data to construct phylogenetic trees, based on the differences in the nucleotide sequences of the virus from different samples, where the branch lengths and relationships in these trees reflect the genetic distances between different viral samples, which in turn can suggest how the virus has spread and evolved over time [Hadfield, 2018; Khare, 2021]. For this purpose, we calculated the correlation between phylogenic distances based on NextStrain data and the strain specific distances from the interaction-based dendrograms obtained from the corresponding clustering for PS, 11G free energy binding and inferred couplings. For this analysis, we kept only the variants with matching NextStrain data (Alpha, Alpha2, Beta, Delta, Delta2, Gamma, Omicron, Omicron BA.2, Omicron BA.4 and Omicron BA.5) (**Supplementary Fig. 2)** and calculated the correlations of the strain distances for all NextStrain-PDB pairs and for each clustering approach (**Figs. 4D, E, F**). Compared to the background distances between all pairs, the distances of the matching pairs exhibited a rather linear relation where PS had the largest correlation (R=0.9) compared to ΔG (R=0.8) and inferred couplings (R=0.6). The strain specific correlations (**Figs. 4G, H, I**), further demonstrates the relation between the ACE2-RBD interactions and the phylogenetic tree where the matching pairs on the diagonal exhibit the highest correlation and all approaches have a block structure discriminating the Omicron group form the other strains. Further inspection shows that the PS-based distance correlates more specifically for related strains like the Alpha and Alpha2 strains and has a stronger separation between the Omicron subvariants and the other strains compared to the ΔG and coupling based distance. Thus, the higher sensitivity of the PS method also allows for a better match between molecular dynamics and phylogenetics.

## Discussion

Virus variants are major drivers of the pandemic and, given changes in transmissibility and disease severeness, it is important to identify VOCs early on. With our MD-based approach, we were able to characterize the observed variants and the relationship between them in a population dynamics independent manner. We were able to connect structural changes in the ACE2-RBD interaction surface to NextStrain phylogenetic data in a timeline based on the emergence of each SARS-CoV-2 variant, conveying understanding of pathogen evolution through space and time, starting from the Alpha variants by the end of 2020, followed by Beta, Gamma, Delta and the Omicron subfamily (**Fig. 1A**).

Based on the structural analysis of the ACE2 and RBD interactive region, we noticed that mutations on residue 501 (N, asparagine to Y, tyrosine) contained in variants such as Alpha, Alpha2, Beta, Gamma and Omicron did not cause major structural deformations in the surrounding residues and tertiary structures, even though such mutation has been shown experimentally to result in one of the highest increases in ACE2 binding affinity conferred by a single RBD mutation [Starr, 2020]. Similar, moderate structural deformations were described for residue 417 (K, lysine to N, asparagine for Beta, Delta, and Omicron; and T, threonine for Gamma). A very strong ACE2 - RBD deformation and consequent loss of contact for residue 484 (E, glutamic acid to K, lysine) was found for variants Alpha2, Beta, Gamma and P2. A similar strong deformation was also observed for the Delta variant with a mutation at a different position (478K) which is only 6 residues away from residue 484. Delta2, which is comprised of a combination of 417N and 478K mutated residues showed much higher flexibility surrounding the mutated area. As for Omicron and its derivative variants, it is very clear that the numerous mutated residues located in this specific area led to an unstable interacting surface area between ACE2 and the RBD.

Overall, our analysis exhibited a strong convergence of structural changes concentrated in the flexible loop area in the interface between ACE2 and RBD for many VOCs **(Sup. Fig. 3, Supplementary PDB Files)**. This result indicates that these shared structural and molecular interaction modifications represent the common biological effect of the VOCs mutations and subsequent epidemiological effects. A recent structural study also identified four key mutations (S477N, G496S, Q498R and N501Y) for the enhanced binding of ACE2 by the Omicron RBD compared to the WT RBD. The effects of the mutations in the RBD for antibody recognition were analyzed, especially for the S371L/S373P/S375F substitutions significantly changing the local conformation of the residing loop to deactivate several class IV neutralizing antibodies [Lan, 2022]. Computational mutagenesis and binding free energies could confirm that the Omicron S protein has a stronger binding to ACE2 than WT SARS-CoV-2, due to significant contributions from residues T478K, Q493K, and Q498R binding energies and doubled electrostatic potential of the RBD-ACE2 complex. Instead of E484K substitution that helped neutralization escape of Beta, Gamma, and Mu variants, Omicron harbors a E484A substitution contributing to a significant drop in the electrostatic potential energies between RBD and mAbs, particularly in Etesevimab, Bamlanivimab, and CT-p59. Mutations in the S protein are prudently devised by the virus that enhances the receptor binding and weakens the mAbs binding to escape the immune response [Shah, 2021].

The normalized RMSF demonstrated common flexible regions throughout the entire ACE2 protein structure (**Fig. 1C**). For the RBD (**Fig. 1D**) we found instabilities in different sections of the protein in dependence on the variants, such as unstable area for the Omicron and Omicron BA.2 variants around residues 370 – 380, while the Omicron BA.3 and Gamma variants have a stringer instability around residues 380 – 400. Around residues 440 – 460 the Beta and Alpha2 variants showed clear RMSF peaks, while around residues 475 – 490 Alpha2 showed a unique 0.2 Å normalized RMSF peak, demonstrating unstable areas unique to these variants. When the RMSF fold change was considered between ACE2 (**Fig. 1E**) and RBD (**Fig. 1F**), it became clear that changes that affected ACE2 were spread over the entire structure, while instability was directed to specific RBD protein segments, and it was variant dependent. Regarding the models and MD simulations used in our study, all the tertiary structures maintained their folding, and simulations were reproducible among replicates (**Supplementary Fig. 1A**). When ACE2 is considered (**Supplementary Fig. 1B**) differences among the variants’ RMSF are negligible, demonstrating ACE2’s stability throughout the simulations and ACE2’s minimal contribution to the structural changes observed. The opposite can be said about the RBD’s RMSF results (**Supplementary Fig. 1C**), most notably residues 360 to 375, where P2 and Alpha demonstrated minimal structural changes, while Gamma and Omicron showed the highest RMSF values. From residue 385 to 395 we observed an area of general structural instability, what would be consistent with this area being comprised of loose loops at the bottom of the RBD structure (for additional information, see **Supplementary Material - Structures**). For Alpha2 and Beta, when residues 440 to 450 were considered, we observed a loop in proximity and displaying several hydrogen bonds between ACE2 chain and the RBD. The residues around 475 to 485 display a mixed behavior depending on the variants, with higher RMSF values for Alpha2, a group of variants with similar behavior to WT (Delta, Delta2, P2 and Gamma), and lower RMSF values than WT (Omicron, Omicron BA.2, Omicron BA.2.12.1, Omicron BA.3, Omicron BA.4, Omicron BA.5, Alpha, and Beta, respectively), possibly indicating a different stability pattern depending on the presence/absence of mutations in the surrounding residues (**Fig. 1E, F**).

Our developed PS approach quantified the residue interactions between ACE2 (**Fig. 2A**) and RBD (**Fig. 2B**) during the simulation time, pinpointing with residue-specific resolution the exact differences in interaction timeframes between specific residues and variants. Our analysis indicates the potential mechanism why Delta mutations can lead to more severe disease [Callaway, 2021] compared to Omicron variants carrying a higher number of mutations and higher infectivity rates [Pulliam, 2021; Grabowski, 2021]. The higher number of mutations might indicate a transition towards an endemic scenario, depending on the interplay of the population’s behavior, demographic structure, susceptibility, and immunity, plus whether viral variants emerge. Different conditions across the world can allow more successful variants to evolve, and these can seed new waves of epidemics. These seeds are tied to a region’s policy decisions and capacity to respond to infections. Even if one region reaches an equilibrium — be that of low or high disease and death — that might be disturbed when a new variant with new characteristics arrives [Katzourakis, 2022]. Overall, this analysis demonstrates that our PS approach can classify mutation-induced changes in virus-host cell binding in a structure-dependent manner and is therefore a powerful tool to monitor and assess the level of concern of newly emerging variants.

The SARS-CoV-2 variants in the PCA results for ACE2 are not clustered by variant and several residues strongly influence the loadings, meaning ACE2 (**Fig. 2C**) is affected by the specific mutations, but only the effects on ACE2 are not enough to enable clear variant classification. The RBD (**Fig. 2D**) is directly affected by the mutations, and we observed the different effects on the PCA loadings in dependence on the variant, making this a great variant classification tool when PS data is applied. Omicron subgroups (Omicron BA.1 and Deltacron, Omicron BA.2 and Omicron BA.2.12.1, Omicron BA.3, Omicron BA.4 and Omicron BA.5) with their high number of mutations are in a completely different spatial area and cluster by themselves separately from all other variants, while still maintain variant-specific resolution that enable the discernment between variants. The clustering of Omicron subvariants in a similar manner could be a positive sign for the future, since other variants such as Beta and Gamma clustered together in our results, and evidence shows that Alpha and Delta variants are more serious than the WT virus in terms of hospitalization, ICU admission, and mortality, as well as Beta and Delta variants, that have a higher risk than the Alpha and Gamma variants [Lin, 2021], whereas Omicron and its derivatives so far appear to be highly contagious but less severe and deadly than the previous variants [Davies, 2022]. For additional insight into PCA results (PC1 to PC5), see **Supplementary Fig. 4**, as well as the PCA loadings in **Supplementary Fig. 5**. To demonstrate that the PS clustering is not biased or related to the chosen methodology, we created mock controls from historical sequences (randomly generated mocks and early SARS-CoV-2 mutations). The results (**Supplementary Fig. 6)** showed similar groupings for the mocks depending on their mutations (weighted, similar positions to SARS-CoV-2 mutations or free, random mutations), following the same separation observed regarding ACE2 (Chain A) and RBD (Chain B) groupings, reassuring the non-bias in our findings.

Regarding Delta clustering closer to P2 and WT than other variants, it has been reported that BLEU452, despite being in the RBD region, does not directly interact with ACE2 [Lan, 2020]. However, BLEU452, together with BPHE490 and BLEU492, forms a hydrophobic patch on the surface of the S protein [Deng, 2021]. A mutation to a highly polar and hydrophilic arginine could potentially introduce local perturbations that could affect how it interacts with a complementary surface. Additionally, BLEU452 is a hotspot located near the negatively charged residues AGLU35, AGLU27 and AASP38 of ACE2. The incorporation of additional charged residues in the vicinity of the binding interface could increase the electrostatic attraction between two proteins. Hence, the mutation of leucine to a positively charged arginine enhances electrostatic complementarity in the interface. Compared to BLEU452, BARG452 was observed to interact more with nearby residues including BSER349, BTYR351, BPHE490, BLEU492 and BSER494. The increased intramolecular interactions could thus increase the stability of the S protein.

When considering the Euclidean distance and similarities between the variants, ACE2 (**Fig. 2E**) seemed to have a more mixed profile of similarities between the variants, while the RBD (**Fig. 2F**) was organized in 2 big groups. One group contained the Omicron subvariants (Omicron BA.1, Omicron BA.2, Omicron BA.2.12.1, Omicron BA.3, Omicron BA.4, Omicron BA.5), Deltacron and Omni, and the other group contained Alpha, Alpha2, Beta, Gamma, Delta, Delta2, P2 and WT. This demonstrates again the similarity between Omicron subvariants versus all other previous variants and the power of the PS to discern between variants and subvariants.

The 11G free energy analysis per residue (energy decomposition) for RBD revealed a considerable energy range from -6 to +6 kcal/mol (**Fig. 3A, Supplementary Fig. 7A, ACE2**) with variants forming both negative and positive energy patches in the heatmap. The resulting signatures allowed for a rough VOC grouping but not for a concise variant classification. The Euclidean distance ΔG heatmap considering the RBD chain showed the cluster with the highest distances for variants P2, Delta and Delta2 when compared to the rest of the variants, as well as a highly mixed variant clustering overall, and thus did not result in a concise variant classification (**Fig. 3B, Supplementary Fig. 7B, ACE2**). The ΔG free energy-based PCA for ACE2 indicated the residues ALYS31, AGLU35, AGLU37, AASP38 and AASP355 as largest separators influencing the sample distribution in this space but with no clear clustering between variants (**Fig. 3C**). For the RBD (**Fig. 3D**) there is a separation between the Omicron subvariants, Deltacron and Omni, the other variants (Alpha, Alpha2, Beta, WT, Delta, Delta2, Gamma,P2). However, unlike the PS PCA, there are no clear subgroups formed.

The complementary approach to infer couplings between putatively interacting SARS-CoV-2 residues by the Thouless-Anderson-Palmer (TAP) approximation for the solution of the inverse Ising problem [Nguyen, 2017], revealed a pattern of strong residue interactions (AGLY354/BGLY502) and a more concise subgrouping of variants into an Omicron cluster (Omicron BA.1, Omicron BA.2, Omicron BA.2.12.1, Omicron BA.3, Omicron BA.4), followed by a cluster containing Omicron BA.5, WT, Omni, Beta, Deltacron, P2; and a group of Delta, Delta2, Gamma, Alpha, Alpha2 (**Fig. 3E**).

In the comparative analysis for the different classification approaches, we investigated the correlation between the phylogenetic NextStrain distances and the distances based on PS, 11G free energy binding, and inferred couplings distances (**Fig. 4A, B, C**). The findings revealed, up to our knowledge, for the first time a strong correlation between the molecular dynamic properties and the phylogenetics. The higher sensitivity of the PS method compared to the ΔG free energy binding and the inferred couplings method led to significant stronger correlations (**Fig. 4D, E, F**). Moreover, the correlation between the strains attested to the superior levels achieved by the PS method, which was consistent with the developments observed during the COVID-19 pandemic (**Fig. 4G, H, I**).

Hence, our PS strategy classifies virus variants into epidemically relevant subgroups, such as distinct Omicron subgroups (Omicron BA.1 and Deltacron, Omicron BA.2 and Omicron BA.2.12.1, Omicron BA.3, Omicron BA.4 and Omicron BA.5), a group containing the P2 and Delta variants, and a larger group containing Alpha, Alpha2, Beta, Gamma, Delta2 variants. The PS variant classification is aligned with findings in terms of the risk of hospitalization, ICU admission, and mortality where the variants Beta and Delta exhibited a higher risk than the Alpha and Gamma variants, and all SARS-COV-2 VOCs have a higher risk of disease severity than the WT virus [Lin, 2021]. Furthermore, Delta infections generated on average 6.2 times more viral RNA copies per milliliter of nasal swabs than Alpha infections during their respective emergence. Our evidence suggests that Delta’s enhanced transmissibility can be attributed to its innate ability to increase infectiousness, but its epidemiological dynamics may vary depending on underlying population attributes [Earnest, 2022]. The German national surveillance data showed e.g., that hospitalization odds associated with Omicron lineage BA.1 or BA.2 infections are up to 80% lower than with Delta infection, primarily in ≥35-year-old. Hospitalized vaccinated Omicron cases’ proportions (2.3% for both lineages) seemed lower than those of the unvaccinated (4.4% for both lineages). Independent of vaccination status, the hospitalization frequency among cases with Delta seemed nearly threefold higher (8.3%) than with Omicron (3.0% for both lineages), suggesting that Omicron inherently causes less severe disease [Sievers, 2022]. The BA.4 and BA.5 subvariants have achieved power from biological changes that allow them to infect more people quickly, possibly due to the spike mutation at position L452R, which was also found in the Delta variant and helps the viral attachment to the human cell. Another vital mutation in BA.4 and BA.5 subvariants is F486V, which occurs in the S protein region close to the attaching site with the human cell, aiding the virus in circumventing the immune system.

With our approach, we were able to classify variants according to epidemic risk, demonstrating that the strain characterization is independent of the population dynamics relying on population sequencing that induces significant delays of two weeks or more but could give early indications for increased transmissibility based on structural and molecular dynamic analyses. Based on the considered synthetic variants Deltacron and Omni that combine mutation from Omicron variants and of either only the Delta or all variants, our analysis suggests that Omicron has been a significant step towards endemics of SARS-CoV-2. The power of the PS method was verified by applying 2 alternative methodologies, 11G free energy binding and inferred couplings between residues, where the PS methodology was superior when considering the ability to differentiate and classify virus variants. Our results suggest that classical affinity estimations, such as ΔG might not capture the full complexity of the virus-receptor interactions, especially in the context of mutations and VOCs, so free energy (ΔG) calculations, while informative, might not adequately represent the dynamic nature of the interactions or the effects of mutations on the virus’s ability to infect and spread. Interestingly, the quantification of interactions by PS and subsequent clustering resembled the phylogenetic difference between the VOCs and thus associates molecular dynamics to phylogenetics for the first time. Overall, our mechanism-based classification is a powerful tool to assess early on the variant-specific epidemic potential which can be integrated in corresponding epidemiological projections and represents therefore an essential element for an early risk assessment of the epidemic dynamics to support political decisions on potential mitigation strategies.

## Author contributions

TA, AH, PM, AT and AS contributed to the conception and design of the study. TA generated all structural models and performed MD simulations. TA and AH performed the analysis. All authors contributed to manuscript revision, read, and approved the submitted version.

