## Supplementary_Figures for "Mechanism-based classification of SARS-CoV-2 Variants by Molecular Dynamics Resembles Phylogenetic Tree"

**Supplementary Figure 1. (A)** RMSD for all SARS-CoV-2 variants. **(B)** RMSF for ACE2. **(C)** RMSD for RBD.

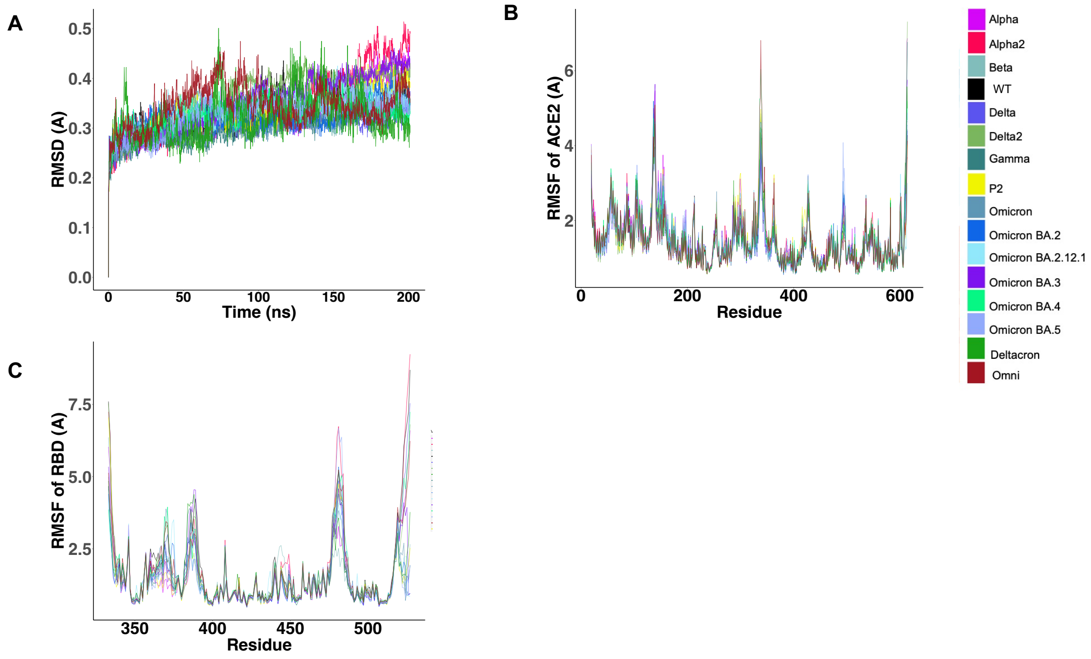

**Supplementary Figure 2.** Variants with matching NexStrain data (Alpha, Alpha2, Beta, Delta, Delta2, Gamma, Omicron, Omicron BA.2, Omicron BA.4 and Omicron BA.5).

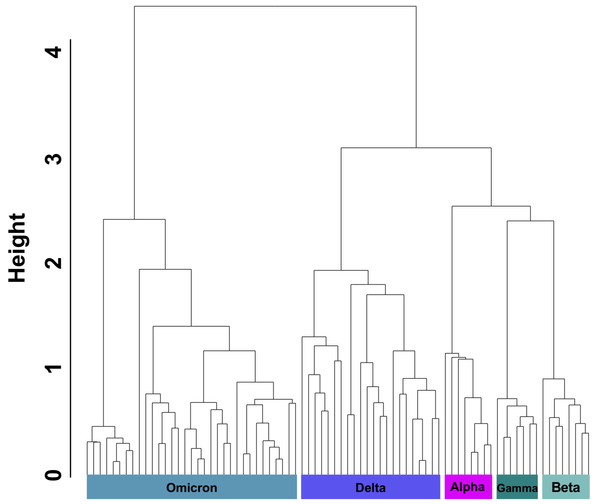

**Supplementary Figure 3.** Our analysis exhibited a strong convergence of structural changes concentrated in the flexible loop area in the interface between ACE2 and RBD for many VOCs, as can be seen in the enumerated residue positions and color-coded structure.

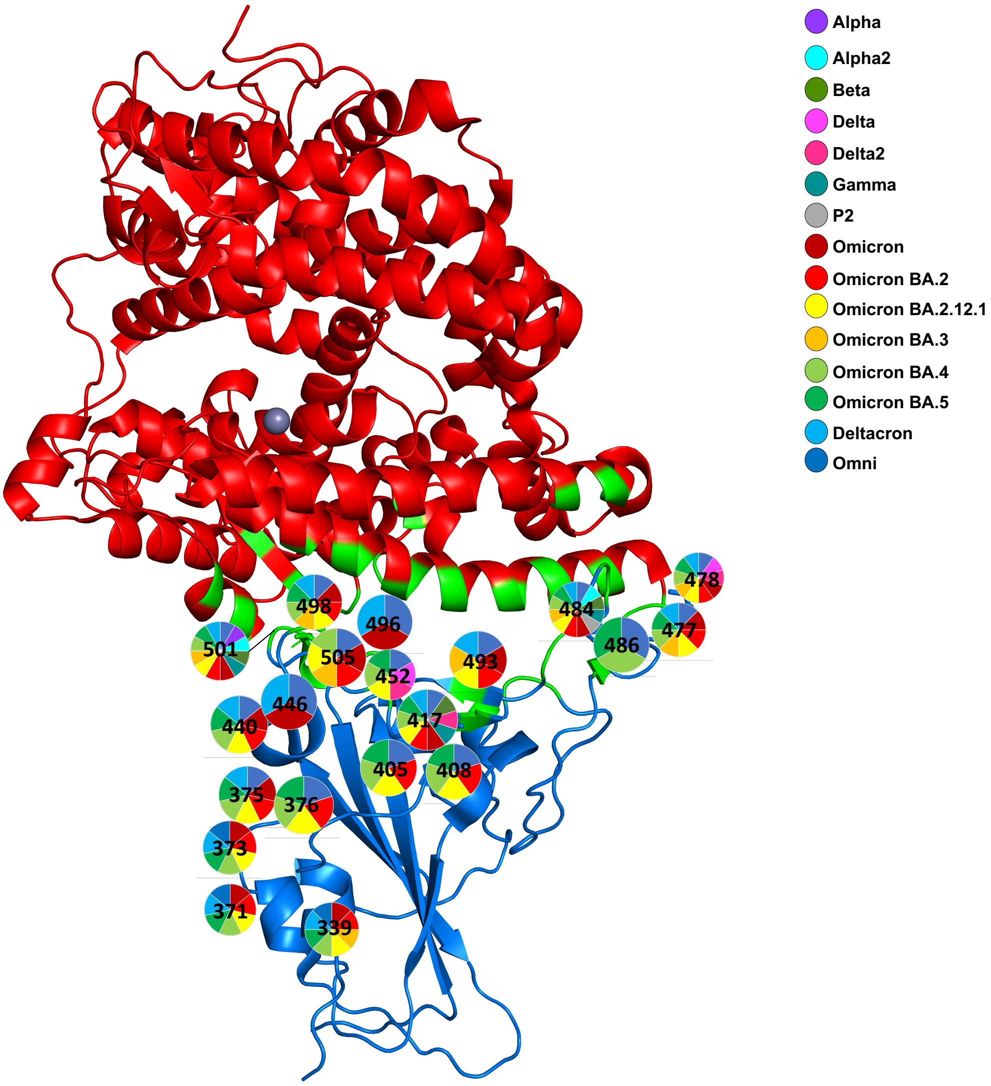

**Supplementary Figure 4.** PCA results (PC1 to PC5) for all variants analyzed.

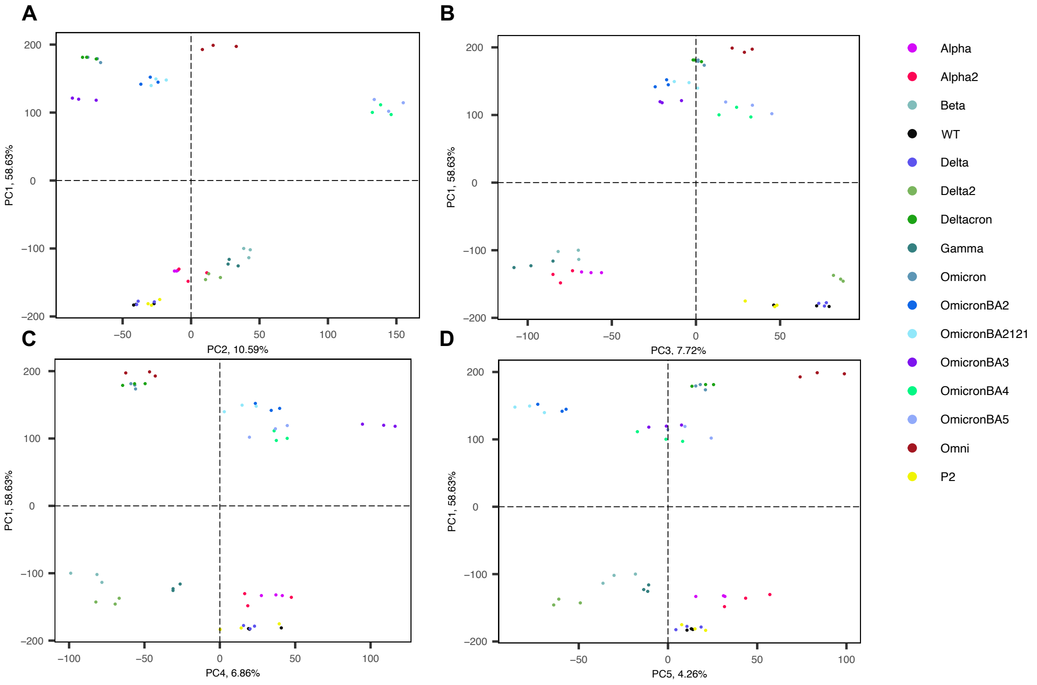


**Supplementary Figure 5.** PCA loadings for PC1 and PC4, where the best separation for P2 and WT was found.

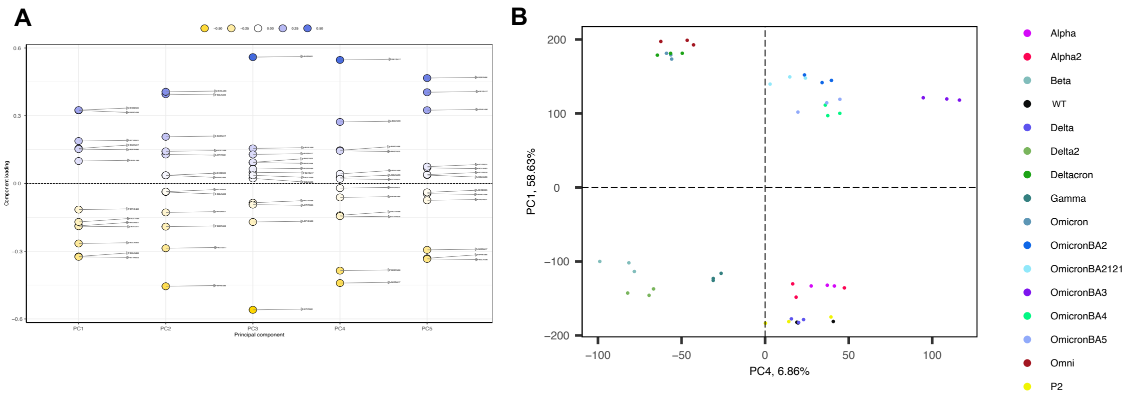

**Supplementary Figure 6.** Mock controls from historical sequences (randomly generated mocks and early SARS-CoV-2 mutations) showed similar groupings for the mocks depending on their mutations (weighted, similar positions to SARS-CoV-2 mutations or free, random mutations), following the same separation observed regarding ACE2 **(A)** and RBD **(B)** groupings, reassuring the non-bias in our findings.

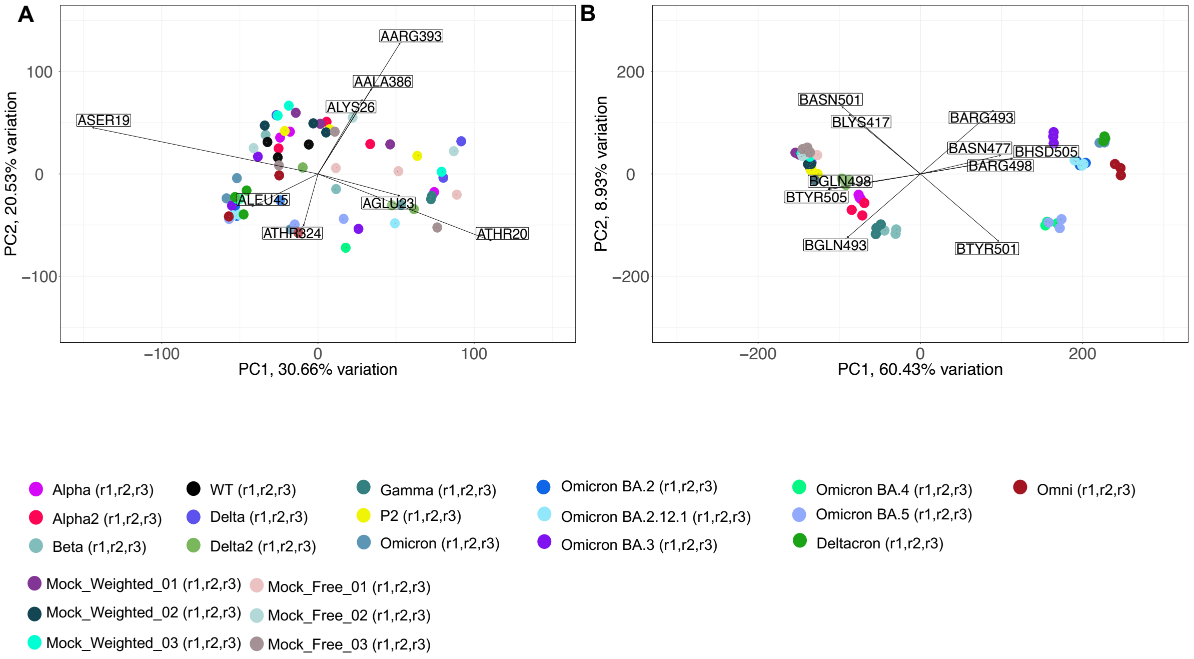

**Supplementary Figure 7. (A)** The ΔG free energy analysis per residue (energy decomposition) for ACE2. **(B)** Euclidean distance ΔG heatmap for ACE2.

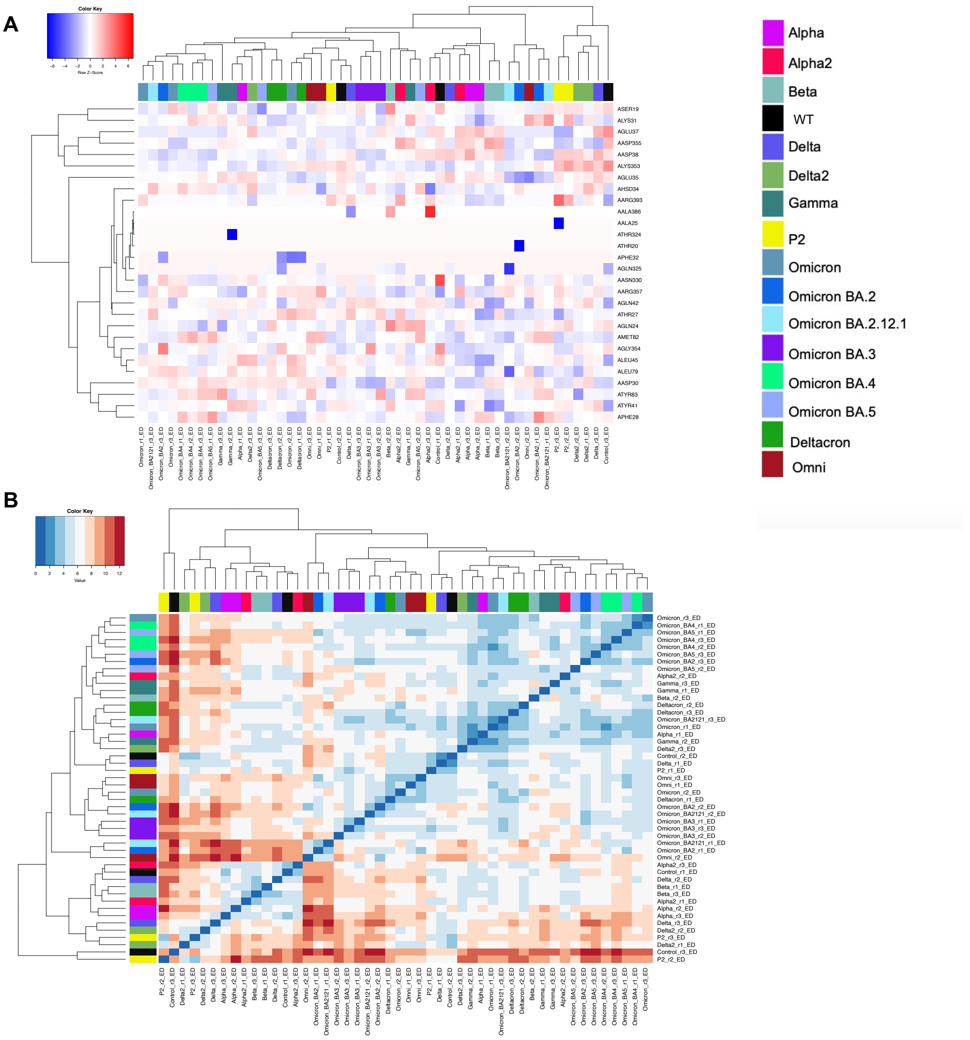


**Supplementary Figure 8.** Groupings for mock controls from historical sequences (randomly generated mocks and early SARS-CoV-2 mutations) when Couplings approach was applied.

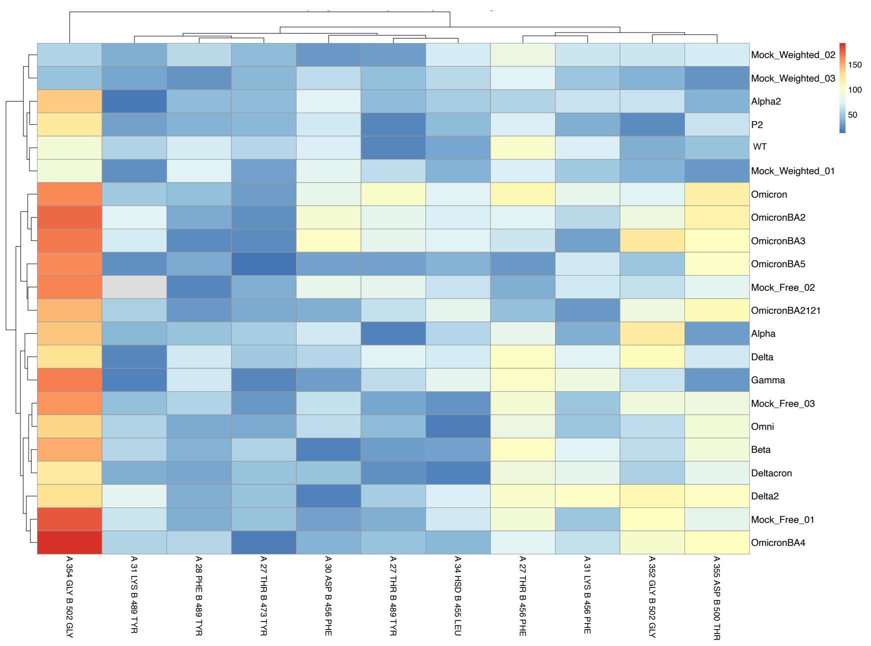
Mock_Weighted_02
Mock_Weighted_03
Alpha2
P2
WT
Mock_Weighted_01
Omicron
OmicronBA2
OmicronBA3
OmicronBA5
Mock_Free_02
OmicronBA2121
Alpha
Delta
Gamma
Mock_Free_03
Omni
Beta
Deltacron
Delta2
Mock_Free_01
OmicronBA4
150
100
50
A 354 GLY B 502 GLY
A 31 LYS B 489 TYR
A 28 PHE B 489 TYR
A 27 THR B 473 TYR
A 30 ASP B 456 PHE
A 27 THR B 489 TYR
A 34 HSD B 455 LEU
A 27 THR B 456 PHE
A 31 LYS B 456 PHE
A 352 GLY B 502 GLY
A 355 ASP B 500 THR
